# Fibers and Glasses are Competing Material States in FUS Protein Condensation

**DOI:** 10.64898/2026.09.28.754945

**Authors:** Liru Feng, Iain Muntz, Ryota Takaki, Lionel Ndamba, Martine Ruer-Gruß, Frank Jülicher, Louise Jawerth

**Affiliations:** Leiden University, Niels Bohrweg 2, 2333 CA Leiden, The Netherlands; Max Planck Institute for the Physics of Complex Systems, Nöthnitzer Str. 38, 01187 Dresden, Germany; Max Planck Institute of Molecular Cell Biology and Genetics, Pfotenhauerstraße 108, 01307 Dresden, Germany

## Abstract

Dense, well-ordered material states of proteins form the amyloid fibers that are a hallmark of neurodegenerative disease in the brain. Beyond forming amyloid fibers, some of these proteins can also adopt other material states termed condensates which are initially liquid-like but evolve to a soft, glassy phase. Fiber growth requires a large supply of monomers and, thus, it is often speculated that fibers emerge from within a dense condensate as it ages and its microscopic dynamics slow into a glassy state. Here, we use the well-established model system Fused in Sarcoma (FUS) to directly observe, quantify and theoretically describe fiber growth and its interplay with condensates. We report the discovery that fibers grow overwhelmingly in the dilute phase surrounding the condensates while the condensates concurrently evolve to a glassy arrested solid. The resulting protein fibers and glassy condensates are both distinct solid-like phases that coexist but do not directly interconvert. Taken together, these findings reveal that there are two competitive aging pathways in FUS condensation that are linked through phase separation kinetics.

## II. BODY

The protein Fused in Sarcoma (FUS) is one of many proteins with intrinsically disordered domains that are implicated in neurodegeneration [1, 2]. These proteins form liquid-like condensates both *in vitro* and in living cells [1, 3–5]. In this process, numerous protein molecules condense from a protein mixture to form a dense condensate phase and a surrounding dilute phase. The liquid phase of FUS has been associated with healthy cellular functions, such as RNA metabolism [6–9] and DNA repair [3, 10]. However, it has also been shown that FUS is prone to form fibers through folding into a cross-beta structure, characteristic of amyloid fibers [11]. Disease-associated mutations of FUS can accelerate fiber formation *in vitro*, suggesting an analogy to the cytoplasmic fibrous inclusions which are the pathological hallmarks of Amyotrophic Lateral Sclerosis (ALS) and fronto-temporal dementia (FTD) [12–15]. In addition to the dilute liquid, dense liquid, and fiber phases, FUS has also been shown to adopt a fourth, glassy material state; it occurs when the condensed liquid phase spontaneously ages through a progressive slowing of its internal dynamics and eventually becoming a soft, glassy solid [16–19]. The interconversion between fibers and various condensate material states is currently unclear; competing hypotheses include that fibers grow rapidly within the high concentration condensates themselves [20–24] and that condensates could act as a metastable sink for fiber formation [25]. However, despite its importance for biological function and disease, the mechanism of FUS fiber growth and its interplay with liquid or glassy condensates has thus far not been directly, experimentally tested. This is crucial for understanding the fundamental processes that govern the role of condensates in biological processes and disease.

In this study we directly observe and quantify fiber growth in the presence of both liquid-like and glassy condensates. Our data and theoretical model support a new paradigm in which fiber growth occurs through the recruitment of material from the dilute phase. Moreover, glassy condensates are a distinct, solid-like material state that is not equivalent to and, in fact competes with, the solid-like fiber material state.

We first study fiber formation in the presence of condensates that are liquid-like. To specifically study fiber growth in the absence of spontaneous nucleation, we use fragmented fibers as sites of nucleation that we refer to as seeds [26]; these seeds rapidly initiate fiber growth (see Methods). We add seeds to a sample containing condensates formed from wild-type FUS tagged with a fluorescent label (FUS-mGFP, *∼*82 kDa) (Fig. 1a and Methods). To each sample, we also add the dye proteostat, which fluoresces when intercalated into cross beta-sheet structures [27]. We can thus observe and discriminate both condensed and fiber phases of FUS using confocal fluorescence microscopy. We observe a three-dimensional volume encompassing several condensates at 20-minute intervals. We define *τ* as the time after the addition of the seeds. At *τ* = 20 minutes, we observe that most seeds are located at the surface of a condensate (Fig. 1b,c and Supplementary Movie 1). At *τ* = 40 minutes, we observe thick fibers emanating from the seed into the dilute phase and a small number of finer fibers growing into the condensate phase (Fig. 1c). Concurrent with this, we observe the growth and thickening of a fibrous shell on the condensate surface (Fig. 1c,d). This shell grows rapidly reaching a thickness of 1-2 *µ*m over 100 minutes, see Fig. 1c,d. At very late times, *τ >* 720 minutes, we observe that condensates have shrunk and obtain crumpled and compacted morphologies (Fig. 1c).

**Figure 1:**
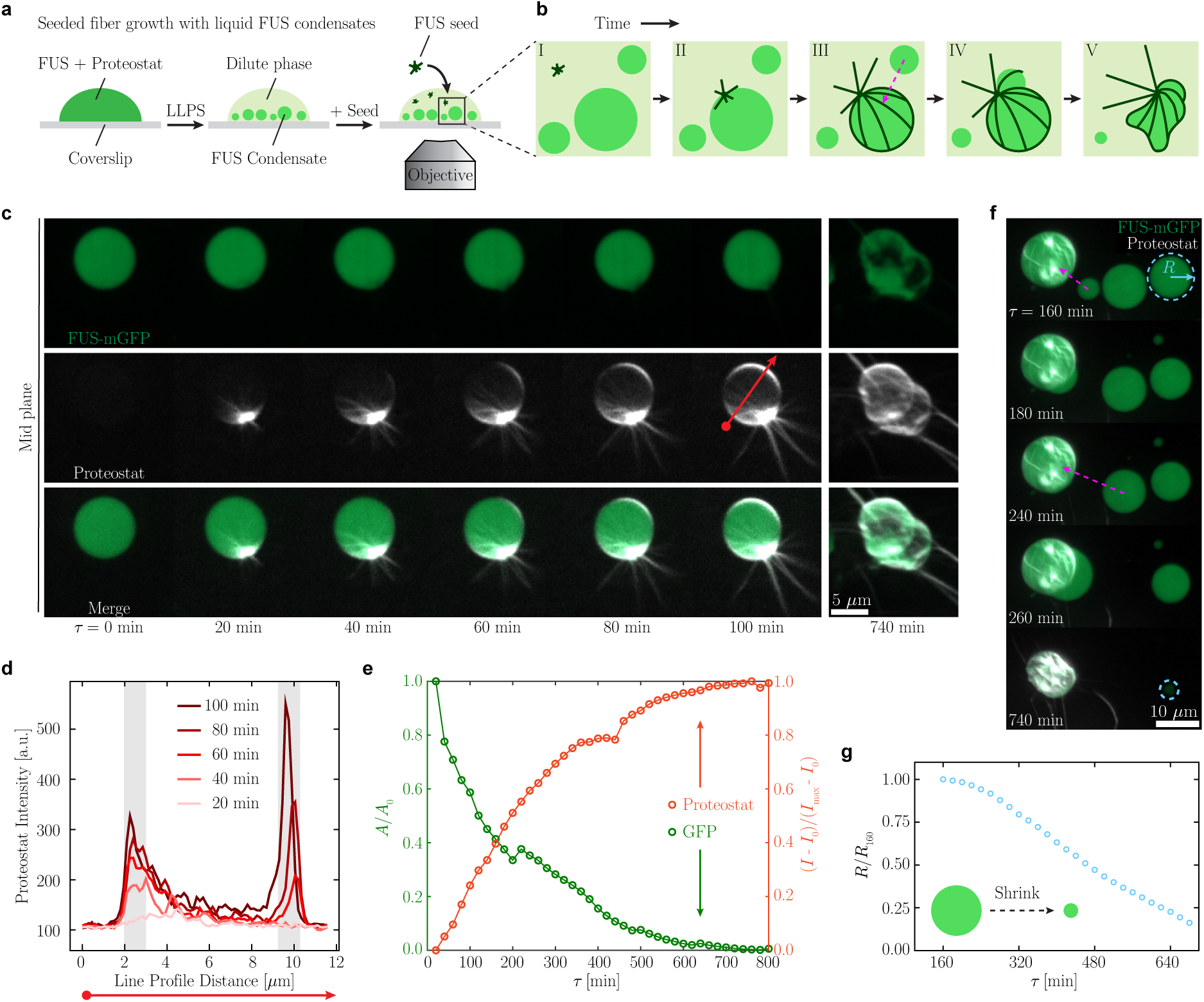
Fibers grow on the surface and in the dilute phase while surrounding condensates shrink and move. a. Liquid-like FUS condensates are reconstituted *in vitro* and the subsequent addition of preformed FUS seeds to the system triggers FUS fiber formation (Supplementary Movie 1). Proteostat, an amyloid-fiber binding dye, is added to localize FUS fibers. b. Mesoscale events during the transition of liquid-like FUS condensates to the fiber phase. Seeds primarily adhere to the surface of FUS condensates (I,II). Fibers form a shell around the condensate surface and grow outward into the dilute phase (II,III). As the fibers continue to elongate, nearby condensates consistently shrink (II-V) and exhibit directed motion toward the fiber growth (magenta arrow in III). In the final stage, condensates and fibers adopt a collapsed morphology (V). c. Confocal images represent the cross-section of a single FUS condensate, green, with an addition of FUS seed. Proteostat which fluoresces within fibers is shown in white. Scale bar, 5 *µ*m. d. Line profile of proteostat fluorescence intensity along the red line shown in c at several time points after the introduction of seeds. e. Total integrated GFP intensity (green) after removing pixels colocalized with proteostat signal and total integrated proteostat intensity (orange) over time after the addition of seeds to FUS condensates. f. Maximum intensity projection of a confocal stack. Shrinking (blue dashed circle) of a surrounding FUS condensate. Dashed magenta arrows illustrate the directed motion of surrounding condensates towards areas of fiber growth. Scale bar, 10 *µ*m. g. The radius of a surrounding FUS condensate, not colocalized with a seed (blue dashed circle in f).

We quantify the fluorescent intensity of the proteostat signal as a proxy for the total fiber formation in the bulk (Fig. 1e orange curve). Concurrently, we determine the total integrated area of GFP signal projected onto a plane, where overlap with proteostat signal has been removed (Fig. 1e green curve). This demonstrates that fibers grow through the recruitment of protein, either directly or indirectly, from the condensed phase.

We also examine condensates that are not in direct contact with a seed or a growing fiber. Surprisingly, we observe numerous dynamic behaviors. In the presence of fiber growth, some condensates will suddenly exhibit directed motion towards a region of high fiber growth. This leads to the appearance that the condensates are being ‘sucked-in’ (Fig. 1f magenta arrows and Supplementary Movie 2). This type of motion may be the result of diffusiophoresis in a concentration gradient of protein in the dilute phase [28, 29]. In addition, we also observe that stationary condensates steadily decrease in size (Fig. 1f dashed blue circle, g and Supplementary Movie 2). This again suggests that the protein concentration surrounding these condensates may be decreasing below the typical dilute phase concentration and dissolving the condensates. By contrast, condensates in a control sample, with an absence of seeds, undergo no directed motion, occasional coalescence and a small degree of shrinking (Extended Data Fig. 2a). Although the dilute phase is often expected to not play a significant role in fiber formation, concentration fluctuations or gradients in this phase could indicate that fiber formation recruits protein directly from the dilute phase altering its concentration.

We therefore tested whether or not fibers are able to form directly from the dilute phase without a condensate precursor. We tested this through two means: first, we nucleated fiber growth from seeds in a buffer containing high salt (300 mM KCl) in which condensates do not form (see Methods and Fig. 2a); second, we added 5-10 *µ*m spherical glass particles to nucleate fiber growth under phase separating conditions. In both cases, we observed the growth of fibers emanating from the seeds or glass particles, respectively, without direct contact to a condensate (Fig. 2a,b). This confirms that fiber growth can occur without a condensate precursor.

**Figure 2:**
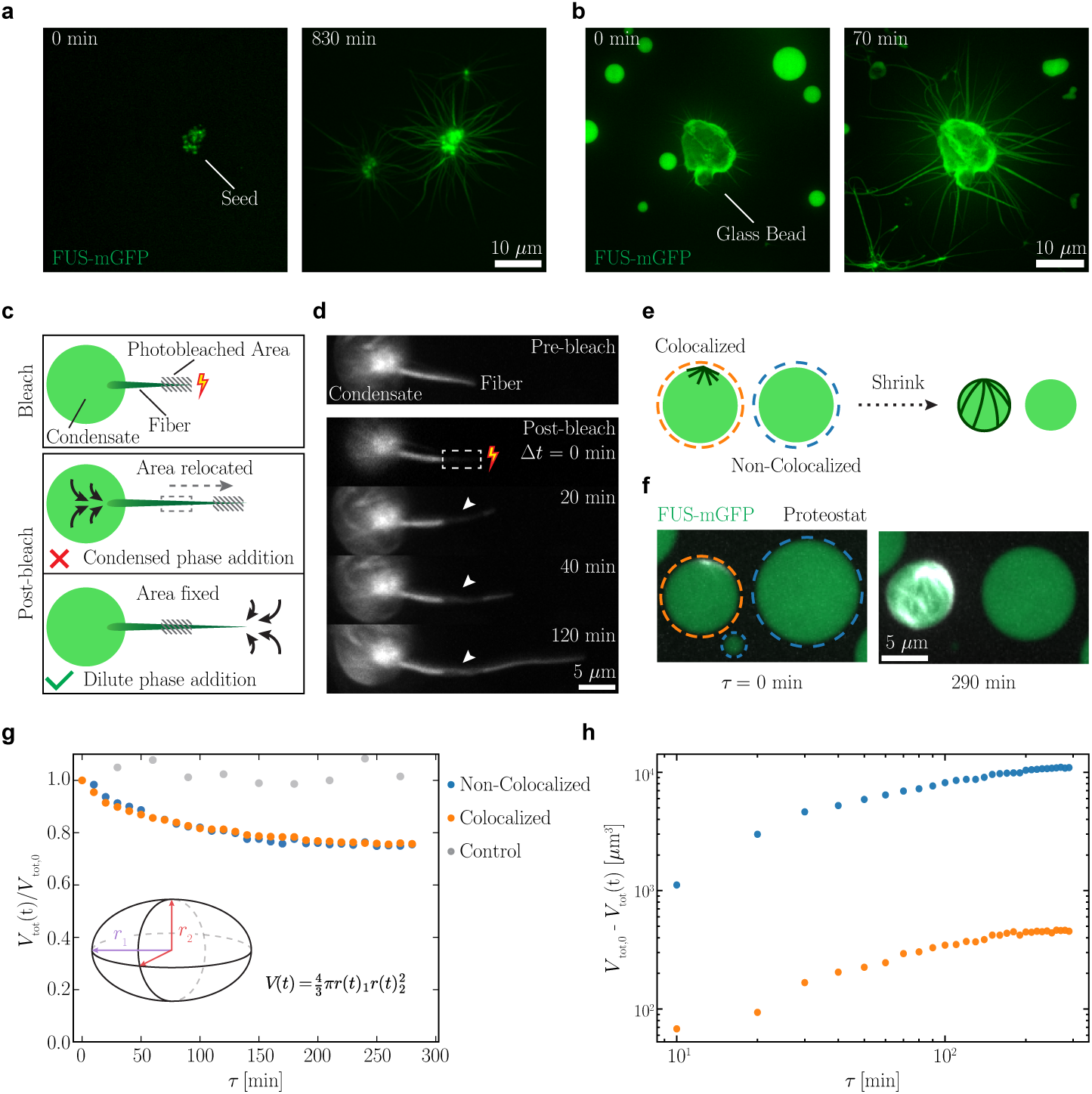
Fibers can and do grow predominantly from dilute phase protein. a. Confocal images of fiber growth from a FUS seed in the absence of FUS condensates in a buffer containing high salt in which condensates do not form. Fibers elongate more than 10 *µ*m from the seed after 830 min. b. Confocal images of fiber growth from a FUS coated glass bead with the presence of liquid-like FUS condensates. Fibers quickly elongate more than 10 *µ*m from the glass bead surface after 1 h. c. Schematic illustration of a FRAP experiment on a growing fiber from a condensate co-localized with a seed. Two possible outcomes are proposed for the location of the bleached area after photobleaching. Either the bleached area translocates indicating growth from the condensed phase or the bleached area remains fixed indicating elongation from the fiber tip. d. Confocal images of a FRAP experiment on a growing fiber from a FUS condensate co-localized with a seed. The bleached area is highlighted by a rectangular dashed box. The bleached area remains fixed as the fiber elongates, indicating that fiber elongation from the tip occurs from protein recruitment in the dilute phase, corresponding to the bottom pictogram of c. e-h. Estimation of total protein fraction recruited from the dilute phase compared to directly from the condensed phase. e-f Schematic illustration and microscopy images of the shrinking FUS condensates over time both co-localized with a seed (orange) or in the absence of a seed (blue). Scale bar, 5 *µ*m. g. Normalized total volumes over time of all condensates co-localized with a seed (orange), not co-localized with a seed (blue), and control (gray; no seed addition). The volume of each condensate (*V_i_*) is measured as an ellipsoid (inset). These show similar trends in condensates both co-localized and not co-localized with a seed. h. Total volume of condensed phase loss over time for condensates co-localized with a seed (orange) and those that are not (blue). The condensates which are not co-localized with a seed lose approximately 24 times the amount of protein compared to the condensates co-localized with a growing seed.

We also tested whether or not a fiber in direct contact with a liquid condensate recruits material preferentially from the dilute or condensed phases. We used a high-intensity laser to photobleach a small area on a fiber growing from a condensate (Fig. 2c,d and Methods, and and Supplementary Movie 4). The photobleached area on the fiber remains permanently dark and can thus act as a fiducial marker. We hypothesized that if the fiber grows recruiting protein from the condensed phase, the fiber would elongate from its condensed phase end and, thus, the bleached area would translocate (Fig. 2c middle panel). By contrast, if material is recruited from the dilute phase the position of the bleached area would remain fixed with respect to the condensate (Fig. 2c lower panel). After photobleaching a fiber in contact with a condensate, we acquired images of the fiber at 20-minute intervals. We observed that the bleached spot remains fixed in its distance from the condensate, consistent with protein recruitment from the dilute phase (Fig. 2d).

We have shown that bleached fibers appear to grow by recruiting molecules from the dilute phase. To investigate if this is indeed the dominant source of monomers for fiber growth, we estimated the amount of material that could be recruited from the condensed phase as compared to that from the dilute phase. We utilize a relevant feature of phase separating proteins: when the dilute phase concentration is lowered below the saturation concentration, condensates will decrease their volume by releasing protein into the dilute phase. [30, 31]. Thus, by carefully quantifying the volume of the condensates not in contact with a fiber we reasoned we could monitor the amount of protein being converted to fibers from the dilute phase. By contrast, when a condensate is in direct contact with a growing fiber, its decrease in volume could be the result of both the direct conversion of protein from the condensed phase to the fiber phase as well as through this indirect conversion from the dilute phase. By comparing the volumes of condensates that colocalize and do not colocalize with growing fibers, we can estimate the fraction of fiber growth which occurs from the dilute phase and directly from the condensed phase. For this analysis, we note that for condensates that are in contact with a growing fiber, their volume loss sets an upper bound on the fraction of protein directly converted from the condensed phase.

We implement the comparison by identifying every condensate in a field of view comprising approximately 150 condensates with a seed density of approximately 1 or 2 per field of view (see Methods and Extended Data Fig. 2b). To estimate condensate volume, we fit an ellipse to each condensate as a function of time and identify both major and minor radii, *r*_1_ and *r*_2_ respectively (Fig. 2g, inset). We measure the total volume of co-localized and not co-localized condensates, respectively, at each time point 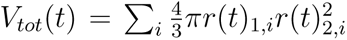 where *i* is an index over each condensate (see Methods). The resulting normalized volume decrease is extremely similar for both condensates that are co-localized with a seed and those that are not, suggesting the loss of condensed phase and its conversion to the fibrous phase is similar in both cases (Fig. 2g). We also compared the total volume of condensed phase loss, *V_tot,_*_0_ *− V_tot_*(*t*), for both condensates colocalized and not co-localized with seeds. The total volume lost is approximately 24 times larger for condensates which are not co-localized with a seed compared to those that are (Fig. 2h blue and orange traces, respectively). This indicates that recruitment of protein from the dilute phase contributes approximately 96% of the protein compared to the condensed phase protein 4% to the total protein converted to fibers under these conditions.

Taken together, our findings suggest growing fibers consume protein predominantly from the dilute phase. As the protein concentration in the dilute phase is lowered, condensates tend to shrink. We developed a theoretical model to test whether this conceptual framework can reproduce our experimental observations. We used an effective droplet model that takes into account the interfacial barrier between the dense and dilute phase for protein diffusion [32–34]. Further, we incorporate the effect of fiber at the system boundary, which acts as a sink for the FUS protein. The core premise of our model is that the shrinking condensates are in a non-equilibrium state [35]. As the dilute phase concentration drops below the saturation concentration, condensate shrinkage is governed by the interfacial conductivity Γ. The interfacial conductivity Γ describes the rate of material flux across the interface and characterizes how readily the material inside the condensate is released to dilute phase. The schematic of our effective droplet model is shown in Fig. 3a. We identify two distinct regimes dictating the condensates’ dynamics depending on the value of Γ.

**Figure 3:**
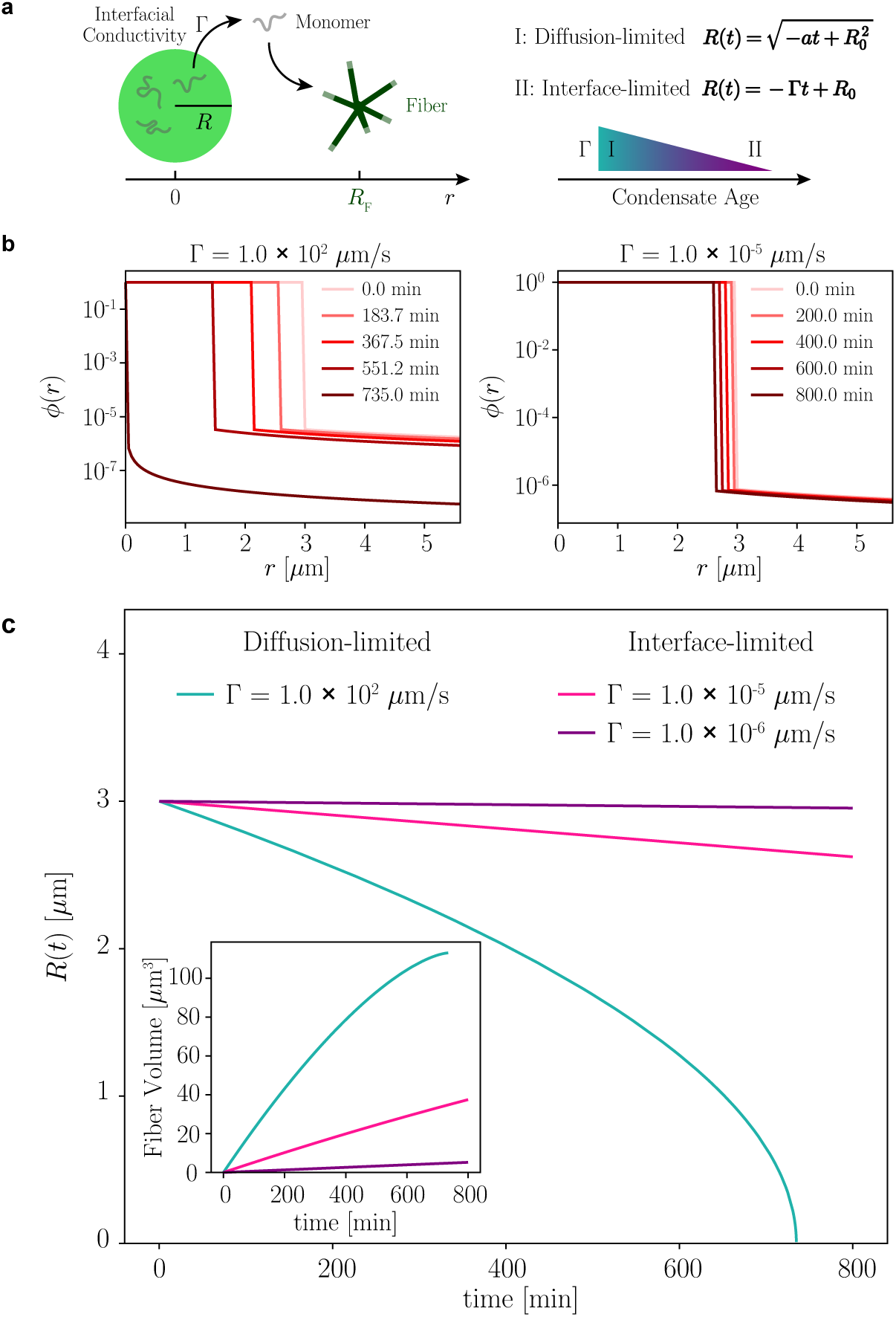
Theoretical model reveals distinct regimes of droplet shrinkage controlled by interfacial conductivity. a. A schematic representation of our effective droplet model. It depicts a spherical condensate of radius *R* (with an initial radius *R*_0_) that indirectly supplies protein for fiber growth occurring at *R_F_*. The rate of material flux across the condensate’s interface is characterized by the conductivity Γ, which decreases as the condensate ages, becoming increasingly glassy. Proteins that leave the condensate then diffuse through the dilute phase before incorporating into the fiber. Depending on the value of Γ, the condensate radius *R* evolves in the diffusion-limited regime (I) or interface-limited regime (II). In the diffusion-limited regime, *a* is a constant determined by the model parameters. b. Simulated protein monomer concentration profiles, *ϕ*(*r*), at five sequential time points for a system in the diffusion-limited regime (left: Γ = 1.0 *×* 10^2^ *µ*m*/*s) and the interface-limited regime (right: Γ = 1.0 *×* 10*^−^*^5^ *µ*m*/*s). c. Condensate radius *R* as a function of time for three different values of Γ. The diffusion-limited regime shows a rapid and nonlinear decrease of condensate radius (Γ = 1.0 *×* 10^2^ *µ*m*/*s). The interface-limited regime shows a slow and linear decrease of condensate radius (Γ = 1.0 *×* 10*^−^*^5^ *µ*m*/*s and Γ = 1.0 *×* 10*^−^*^6^ *µ*m*/*s). The inset shows the corresponding accumulated fiber volume over time, demonstrating that fiber formation is significantly enhanced in the diffusion-limited regime.

The first regime is the diffusion-limited regime, which occurs when interfacial transport is fast compared to bulk diffusion (i.e., when Γ *≫ D*_out_*/R*, where *D*_out_ is the external diffusion coefficient and *R* is the condensate radius). In this limit, quasi-equilibrium conditions at the interface are recovered, and a condensate shrinks with the functional form: 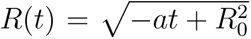, where is a constant determined by the model parameters (see Methods).

The second regime, the interfacial-limited regime, dominates in the opposite limit (Γ *≪ D*_out_*/R*). Here, slow kinetics at the interface become the bottleneck for shrinkage, and non-equilibrium effects are prominent. This regime is characterized by a linear decrease in the condensate radius: *R*(*t*) = *−*Γ*t* + *R*_0_.

For the diffusion-limited regime (Γ = 1.0 *×* 10^2^ *µ*m/s), the condensate exhibits more rapid shrinking compared to the interfacial-limited regime (Γ = 1.0 *×* 10*^−^*^5^ *µ*m/s), see Fig. 3b. These distinct shrinkage kinetics are further illustrated in Fig. 3c. Importantly, Γ controls the rate at which FUS monomers are replenished in the external phase. A higher conductivity sustains a larger concentration gradient towards the fiber sink, thereby enhancing the rate of fiber formation, as quantified in the inset of Fig. 3c.

An interesting consequence of this model is that if the protein in the condensed phase cannot be rapidly released to the dilute phase, fiber formation should be slowed. This would be the case if the condensed phase is no longer liquid-like but, instead, adopts a glassy state in which protein is likely to be jammed or sequestered. Although consistent with our conceptual understanding and theroetical model, this conjecture is contrary to the common assumption that a glassy phase promotes fiber formation. We tested this conjecture experimentally. We repeated our experiment and nucleated fiber formation in the presence of FUS condensates with varying degrees of glassiness. We have previously shown that glass-like aging of condensates occurs spontaneously over time after initial condensate formation [16]. We therefore formed 3 identical samples of FUS condensates in the absence of seeds and let them incubate for *t_w_* = 1 h, *t_w_* = 24 h, and *t_w_* = 48 h, respectively, where the waiting time, *t_w_*, is defined as the incubation time (Fig. 4a). We used fluo-rescence recovery after photobleaching (FRAP) as well as dissolution to confirm that the samples evolve to an increasingly arrested, glassy state during incubation (Extended Data Fig. 3). After the respective incubation time, we added numerous fiber seeds to nucleate rapid fiber growth in each sample (Fig. 4a).

**Figure 4:**
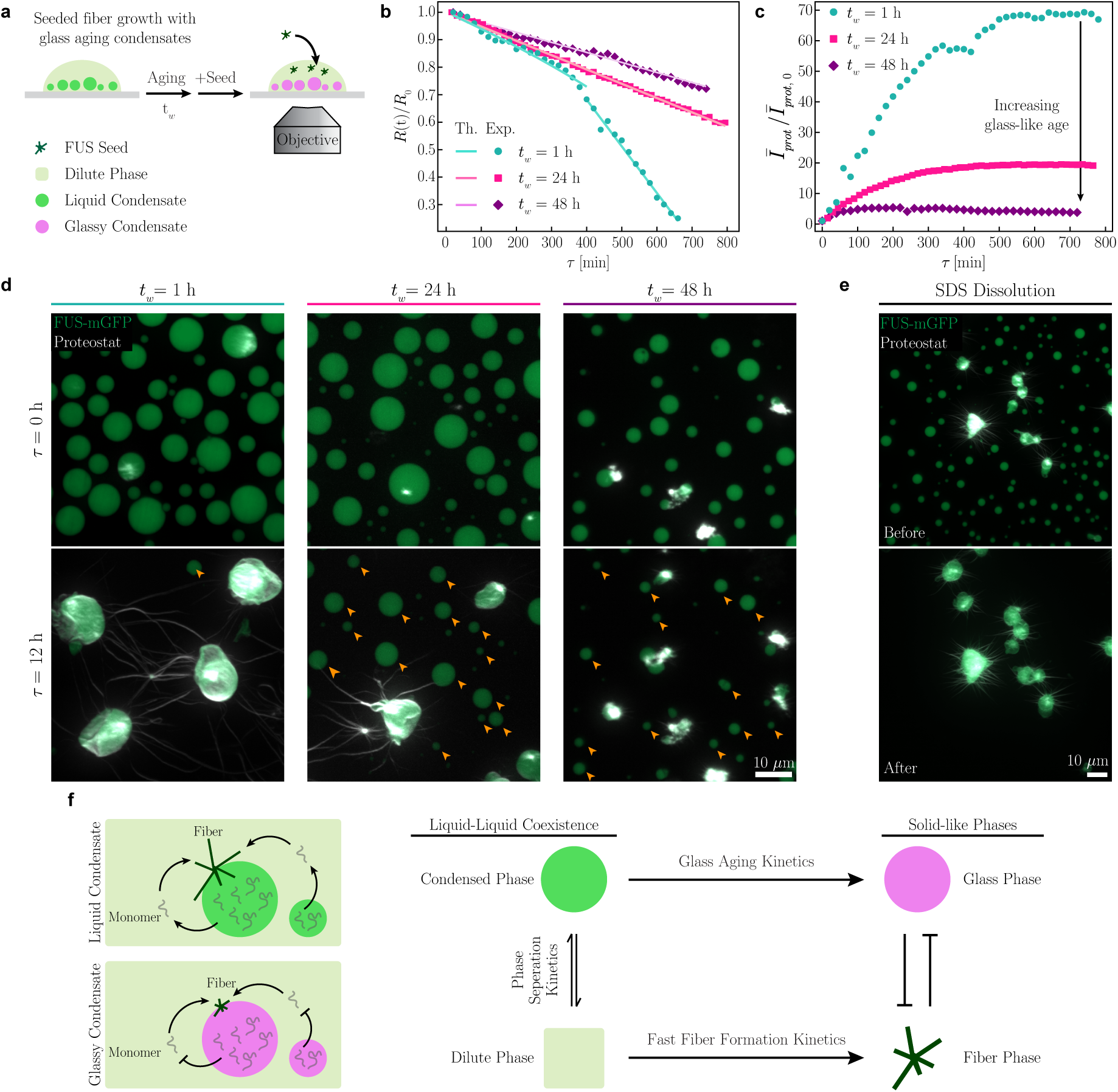
Condensate glassiness is a distinct solid state that hinders fiber formation. a. Liquid-like FUS condensates are reconstituted and incubated for varying durations (*t_w_*). The subsequent addition of seeds triggers fiber formation. The incubation time *t_w_*, following phase separation, determines the extent of condensate glassiness. Longer incubation times result in more condensate material remaining in a glassy state. The condensate glassiness has been demonstrated in previous work [16] and is examined by our FRAP experiment and high-salt dissolution assay (Extended Data Fig. 3) b-d. The experiment of seeding after aging FUS condensates. b. Normalized condensate radius, *R*(*t*)*/R*_0_, after seed addition. The lines represent fits to the data using either the interface-limited (*R ∝ t*) or diffusion-limited (*R*^2^ *∝ t*) expressions. The interface-limited expression accurately describes the data for *t_w_* = 24 h and *t_w_* = 48 h, yielding Γ values of 1.2 *×* 10*^−^*^3^ *µ*m*/*s and 5.5 *×* 10*^−^*^4^ *µ*m*/*s, respectively. The *t_w_* = 1 h data exhibits a crossover, initially following the diffusion-limited behavior (*τ <* 400 min, *a* = 5.9 *×* 10*^−^*^3^ *µ*m^2^*/*s) before transitioning to an interface-limited behavior at later times (*τ >* 400 min, Γ = 3.8 *×* 10*^−^*^3^ *µ*m*/*s). c. Normalized fluorescence intensity over time for proteostat indicating slowed fiber growth for glassier condensates. The normalized fluorescence intensities plateau at 68.2 *±* 0.7, 19.39 *±* 0.11, and 5.2 *±* 0.2 for *t_w_* = 1 h, *t_w_* = 24 h, and *t_w_* = 48 h respectively. While the initial gradients are 0.21 *±* 0.05 s*^−^*^1^, 0.096 *±* 0.005 s*^−^*^1^, and 0.059 *±* 0.009 s*^−^*^1^ for *t_w_* = 1 h, *t_w_* = 24 h, and *t_w_* = 48 h respectively. d. Maximum intensity projection confocal images of FUS liquid-to-fiber transition induced by seeding with aging condensates (see Supplementary Movie 5-7). Condensates maintained over the course of 12 hours after seed addition are highlighted with yellow arrowheads. Scale bar, 10 *µ*m. e. Glassy condensates and fibers coexist after FUS condensates are aged for 24 hours and incubated with seeds for an additional 19 hours. After SDS treatment, FUS fibers resist the detergent, whereas glassy FUS condensates fully dissolve. Scale bar, 10 *µ*m. f. Pictogram of fiber growth in the presence of condensates. (left) Liquid-like FUS condensate acts as a reservoir to fuel fiber growth, while glass-like aging of FUS condensate slows fiber growth from seeds. (right) Liquid-like condensed and dilute phases are linked but each evolves distinctly to a solid-like phase: The condensates evolve to a solid-like, glassy state with slow dynamics while the dilute phase evolves to solid-like, fibers.

We observed a three-dimensional region encompassing numerous condensates at 20-minute intervals over 12 hours using confocal fluorescence microscopy. In all cases, fiber growth occurred at the seeds after 20 minutes. For a sample with a 1-hour incubation time, we reproduced the behavior reported in Fig. 1-3 with an almost total loss of condensed phase protein after 12 hours with significant fiber growth (Fig. 4b,c cyan traces, 4d, and Supplementary Movie 5). By contrast, for the sample incubated 24 hours, we observed a reduced quantity of fibers with significant condensed phase remaining after 12 hours (Fig. 4d yellow arrows, and Supplementary Movie 6). This qualitative effect was more pronounced in the case of condensates with 48 hours of incubation with even fewer apparent fibers and numerous remaining condensates after 12 hours (Fig. 4b,c purple traces, and Supplementary Movie 7).

We quantified these observations and fit them with our theoretical expectations. For condensates incubated for *t_w_* = 1 h, we determine that the condensates initially shrink with a radius, *R*^2^ *∝ t* consistent with a diffusion-limited regime. At later times, *τ >* 400 min, the sample progressively ages and the radius scales instead with *R ∝ t* indicating a transition to the interface-limited regime; the data are well fit with Γ = 3.8 *·* 10*^−^*^3^*µ*m*/*s (see Fig. 4b cyan markers). This can be understood as the liquid-like condensates with little incubation time, *t_w_* = 1 h, release their material quickly into the dilute phase, initially, but subsequently transition to a jammed state. This is suggestive of glassy aging occurring *in situ* during the course of our 12-hour observation. For condensates incubated for *t_w_* = 24 h and *t_w_* = 48 h, respectively, the shrinking rate further decreases with incubation time and is well-fit with interface-limited behavior, Γ = 1.2 *·* 10*^−^*^3^*µ*m*/*s and Γ = 0.55 *·* 10*^−^*^3^*µ*m*/*s, respectively (see Fig. 4b pink and purple markers). The steady decrease in interfacial conductivity, Γ, with condensate age, *t_w_* suggests that the protein is increasingly jammed/sequestered consistent with our expectations of an aging glass.

Lastly, we tested whether increased condensate age, *t_w_* does, indeed, decrease the total amount of fiber formation. We quantified the normalized amount of fiber growth during each experiment using the fluorescence intensity from the proteostat dye scaled to the initial intensity of the exogenously added seeds, see Methods. Samples from all 3 incubation times, *t_w_*, exhibit a rapid increase in fiber amount at early times followed by a plateau at long times. Consistent with our expectation, the initial rate of normalized fiber growth decreases with increasing condensate waiting time, 0.21 s*^−^*^1^, 0.096 s*^−^*^1^ and 0.059 s*^−^*^1^, for *t_w_* = 1 h, *t_w_* = 24 h, and *t_w_* = 48 h, respectively, (see Fig. 4c and Methods). The final fiber plateau value also decreases with increased waiting time; we determined 68, 19, and 5, for the samples with *t_w_* = 1 h, *t_w_* = 24 h, and *t_w_* = 48 h, respectively. (see Fig. 4c and Methods). From these data, we conclude that the protein in the glassy condensates is sequestered and released only very slowly into the dilute phase solution (Fig. 4d), thus decreasing fiber formation.

These data suggest that the incubated condensates are in a glassy phase that is distinct from the fiber phase of protein. To test this, we added a sodium dodecyl sulfate (SDS) solution (10% w/v SDS) to a sample which had been incubated for 24 hours without seeds and subsequently incubated with seeds for an additional 19 hours. Amyloid fibers characteristically do not dissolve with SDS and this is therefore a common test to discriminate amyloid fibers from other protein states [36]. Upon SDS treatment, the glassy condensates completely dissolved within 2 hours while the fibers persisted (Fig. 4e). This confirms that the glassy, condensed phase although arrested is distinct from the fiber phase of protein.

## III. CONCLUSION AND DISCUSSION

In this study, we directly observed and quantified the growth of FUS fibers and their interplay with both liquid and glassy condensates. We discovered that FUS fibers grow primarily through the recruitment of protein from the low-concentration, dilute phase that surrounds condensates rather than within the condensate interior. Concurrent to dilute phase fiber growth, the condensates themselves evolve into soft, glassy solids. The condensates are, however, not entirely passive but contribute to fiber growth by releasing protein to maintain the dilute phase concentration. Theoretical modeling reveals that at early times, this release is limited by diffusion of monomers in the dilute phase but becomes interface-limited as the condensates become increasingly solid-like. Moreover, our data show that while both are solid-like, fibers and glassy condensates are two distinct material states. Taken together, we arrive at a new paradigm for FUS phase separation and fiber growth: the dilute phase and the condensed phase each age through separate processes to form distinct solid-like phases that do not directly inter-convert; however, at all times they remain connected through the local kinetics of phase separation, see Fig. 4f. Thus, the glassy solid condensate is not a precursor to fiber growth and, in fact, impedes sustained fiber growth.

A recent, related study has also suggested that that fibers grow in the dilute phase of a phase separated protein system. The authors estimated the free energy differences between fibers and condensates based on dilute phase concentrations in the presence of fibers and without, using equilibrium considerations [25]. However, our work uncovers that the system is inherently non-equilibrium: condensates themselves undergo glass-like aging and are out-of-equilibrium. This becomes even more apparent when fibers grow: the dilute phase concentration likely lowers far below the saturation concentration of condensates, which slowly dissolve in a process governed by surface conductivity. Thus, a theoretical framework that incorporates the system’s non-equilibrium features, as we propose here, is essential for a description of the interplay of condensates and fibers. In summary, we discover that glass-like aging of condensates and amyloid fiber formation can act as competing mechanisms within the FUS condensate system. As amyloid fiber formation is often associated with disease and biological dysfunction, an exciting avenue of future research will be to identify the role of glass-like aging in more complex phase separating systems including those found within living systems.

## IV. ACKNOWLEDGMENTS

We would like to thank Wilson Poon, Tyler Harmon, Lars Hubatsch, Anthony Hyman and L. Mahadevan for insightful discussions; Régis Lemaitre and Anthony Hyman for assistance with protein production; and Anna Bakker for preliminary experiments.

