## Supplementary Information for "Fibers and Glasses are Competing Material States in FUS Protein Condensation"

### I. MATERIALS AND EXPERIMENTAL METHODS

#### A. Protein Expression and Purification

FUS-mGFP (82.43 kDa) was expressed and purified as described previously [3,16]. FUS-mGFP was subcloned into a construct encoding MBP-3C-FUS-mGFP-3C-6xHis. In this construct, 3C is the PreScission protease cleavage site and mGFP is a monomeric GFP

bearing an A206K mutation [37]. This construct was compatible with the FlexiBAC insect cell system [38] and the viruses containing the protein construct were generated with the FlexiBAC system. The MBP-3C-FUS-mGFP-3C-6xHis protein was expressed in a Sf9 insect cells (Expression Systems) and harvested 72 hours post-infection. For purifying FUS-mGFP, MBP-3C-FUS-mGFP-3C-6xHis, the cell pellet was collected by spinning the cells at room temperature (23 °C). The pellet was resuspended and the resuspended cells were lysed by sonication and clarified by centrifugation at 25,000 rpm for 1 hour. The supernatant was collected and filtered with a bottle top vacuum filter (Nalgene, Thermo Scientific) with a pore size of 0.45 μm filter membrane. The filtered supernatant was loaded onto Ni-NTA FPLC columns (Protino, Macherey-Nagel). The columns were pre-equilibrated with the Ni-NTA binding buffer containing 50 mM HEPES pH 7.3, 1 M KCl, 5% Glycerol, 1 mM DTT, and 10 mM Imidazole. The columns were washed by 10 column volume Ni-NTA binding buffer. The MBP-3C-FUS-mGFP-3C-6xHis protein was eluted by the elution buffer containing 50 mM HEPES pH 7.3, 1 M KCl, 5% Glycerol, 1 mM DTT, and 250 mM Imidazole. The His-tag purified MBP-3C-FUS-mGFP-3C-6xHis protein was loaded onto Amylose gravity flow columns (New England Biolabs). The columns were pre-equilibrated with the Amylose binding buffer containing 50 mM HEPES pH 7.3, 500 mM KCl, 5% Glycerol, and 1 mM DTT. The columns were washed by 10 column volume binding buffer containing 50 mM HEPES pH 7.3, 1 M KCl, 5% Glycerol, and 1 mM DTT. The MBP-3C-FUS-mGFP-3C-6xHis protein was eluted by the elution buffer containing 50 mM HEPES pH 7.3, 1 M KCl, 5% Glycerol, 1 mM DTT, and 250 mM maltose. The MBP-tag purified protein solution was concentrated to 5 mL. The Pierce HRV 3C protease (Thermo Scientific) was added to the concentrated MBP-tag purified protein at a 1:50 ratio. After 4 hours cleavage, the protein solution containing the released MBP, FUS-mGFP, 6xHis, and PreScission protease was filtered and loaded onto the gel-filtration column (16/600 Superdex 200 pg, Cytiva) that was pre-equilibrated with the storage buffer containing 50 mM HEPES pH 7.3, 500 mM KCl, 1 mM DTT, and 5% Glycerol. The protein mixture was further purified over the gel-filtration chromatography and the peak fractions containing pure FUS-mGFP were pooled, concentrated by Amicon Ultra centrifugal filters (30 kDa MWCO; Millipore), and aliquoted in PCR tubes (STARLAB). FUS-mGFP stock was flash-frozen by liquid nitrogen and stored at -80 °C. Purity of the protein was confirmed by SDS-PAGE. Concentration of the protein was measured at 280 nm with Microvolume UV/Vis Spectrophotometers (Implen).

### B. FUS Seed Preparation

FUS stock (90 $\mu\text{M}$ FUS, 50 mM HEPES pH 7.3, 1 mM DTT, 500 mM KCl) was incubated quiescently in PCR tubes at room temperature (23 °C) for 7 days. FUS fiber fragmentation was achieved by sonication. For large seeds ($\sim 5\ \mu\text{m}$ size), the fiber solution was sonicated for 1 hour using an ultrasonic water bath at room temperature. For small seeds ($\sim 1\ \mu\text{m}$ size), the fiber solution was sonicated for 10 minutes using a sonifier (Branson SFX150) with a 3 mm microtip, at 20% amplitude in consecutive 5-s on/off cycles on an ice bath. Fragmented FUS fibers (i.e. seeds) were aliquoted into protein lobind tubes (Eppendorf) and flash-frozen in liquid nitrogen and stored at -80 °C. The concentration of FUS seed was estimated based on the monomeric FUS concentration.

### C. Assay for Seeded Fiber Growth in the Presence of Condensates

FUS condensation is initiated through dilution of FUS stock with a buffer that contains no KCl (dilution buffer, 50 mM HEPES, 1 mM DTT at pH 7.3) along with proteostat dye stock (Enzo Life Sciences) to a final FUS concentration of 15-17 $\mu\text{M}$ FUS, salt concentration of 100 mM KCl, and a 1000-fold dilution of proteostat. The FUS samples were added into the 384-well ULA-coated microplates (PhenoPlate, Revvity). Samples were aged quiescently at room temperature (23 °C) for different durations, $t_w$, of 1 h, 24 h, and 48 h. The FUS seed stock was diluted with dilution buffer to various seed concentrations and subsequently added to fresh or aged FUS condensates. The FUS seed concentrations varied between 300-700 nM for most samples. For the samples in which we analyze the respective fraction of protein arising from the dilute phase or the condensed phase (depicted in Fig. 2), we attempt to find conditions in which we have approximately 1 or 2 seeds per field of view which corresponded to approximately 20 nM.

Note: We have also tested other red amyloid-binding dyes, such as Amytracker 680 (Ebba Biotech), Congo Red (Sigma), and CRANAD-2 (Sigma). However, in our preliminary tests, we found that Congo Red dissolved the condensates, Amytracker caused dye aggregates within the condensates, and CRANAD-2 failed to stain fibers. In contrast, the presence of proteostat does not alter FUS fiber formation or phase separation.

##### **D. Assay for Seeded FUS Fiber Growth in the Absence of FUS Condensates**

FUS stock was diluted to a salt concentration of 300 mM KCl and a FUS concentration of 15 μM, similar to the concentration used for initiating FUS LLPS. FUS seed stock was diluted with dilution buffer to various seed concentrations, with 300 mM KCl prior to addition to FUS samples to induce seeding. The final condition illustrated in Fig. 2a is 300 mM KCl, 15 μM FUS, 0.7 μM seed, 50 mM HEPES pH 7.3, and 1 mM DTT. Fiber growth was imaged using confocal microscopy, as described in section I F.

##### **E. Assay for Seeding FUS Fiber Growth from Glass Beads**

A solution of glass beads (Sigma; 1.5% w/v) was incubated in FUS solution (15 μM FUS, 50 mM HEPES pH 7.3, 1 mM DTT, 400 mM KCl) in a protein lobind tubes (Eppendorf) quiescently for a week at room temperature (23 °C). The coated glass beads were subsequently washed 3 times with dilution buffer (50 mM HEPES, 1 mM DTT, pH 7.3) using centrifugation to separate the particles from the supernatant. The washed glass beads were then diluted 10 times and stored in dilution buffer. Finally, glass beads were added to a condensate sample with a final concentration of 4.5 μM FUS in a buffer of 50 mM HEPES pH 7.3, 1 mM DTT, and different concentrations of KCl. These were observed with confocal microscopy (Fig. 2b, 100 mM KCl, with LLPS; Extended Data Fig. 1b, 200 mM KCl, without LLPS).

##### **F. Confocal Fluorescence Microscopy**

The FUS samples were imaged with a spinning disk confocal (Yokogawa CSU-X1) on an inverted microscope (Nikon Ti-E) equipped with a water-immersion 60× Plan Fluor NA 1.27 objective lens, a PiezoZ drive (MCL NanoDrive), and the Perfect Focus system. The microscope was mounted on a vibration-isolation air table. The pixel size for the 60× objective is 0.15 μm. FUS-mGFP and proteostat were excited with a 488 nm and a 561 nm solid-state laser. To image FUS-mGFP and proteostat simultaneously, we illuminated the sample sequentially using excitation wavelengths of 488 nm and 561 nm. The two fluorescent signals were allocated to the same image sensor through a multiband emission filter, producing two distinct images. The exposure time for each frame was 50 ms. Z-stacks were taken with a step size of $\sim 200$ nm, with the shutter closed in-between steps. Images were obtained with a sCMOS camera (Prime 95B and Andor Zyla) controlled with Nikon NIS-elements AR software (version 5.42.04).

#### G. Sodium Dodecyl Sulfate Dissolution Assay

A sodium dodecyl sulfate (SDS) protein denaturant solution (10% w/v), at 5 times the sample volume, was added to the sample well containing FUS condensates which had been prepared as described in section IC with an incubation time of 24 hours without seeds followed by an additional 19 hours with seeds at room temperature (23 °C). Confocal fluorescence images were taken before SDS addition and 45 minutes after the addition.

#### H. Fluorescence Recovery after Photobleaching

Fluorescence recovery after photobleaching (FRAP) experiments were performed using a confocal microscope using the FRAPPA photomanipulation system. To test fiber growth from a fiber in contact with a condensate, the fiber tip was bleached by focusing a high-intensity 488 nm laser beam on a rectangular area (Fig. 2d) for a dwell time of 1000 ms. Recovery was monitored for 12 minutes following bleaching.

In the whole FRAP experiment shown in Extended Data Fig. 3a, condensates aged for different time periods (1 hour, 24 hours, and 48 hours) at room temperature (23 °C) were bleached by focusing a high-intensity 488 nm laser on a circular area with a diameter equal to that of the condensate, with a 1000 ms dwell time. FRAP images are acquired at a rate of 2 seconds. Pre-bleach images are acquired for 2 seconds. After bleaching, the mean fluorescence intensity of the bleach area is around 10% of the pre-bleach intensity. Fluorescence recovery was monitored for around 5 minutes. Small translations in the image were corrected through image registration implemented using FIJI (National Institutes of Health, Bethesda, MD) and the StackReg Plugin with the Translation option. FRAP curves were analyzed using custom scripts in Python. The intensity at the first post-bleach time point ($I_{\text{post}}$) was subtracted from the intensity of the bleached region, $I(t)$, and the result was scaled by the difference between the pre-bleach intensity ($I_{\text{pre}}$) and $I_{\text{post}}$,

$$I_{\text{norm}} = \frac{I(t) - I_{\text{post}}}{I_{\text{pre}} - I_{\text{post}}} \quad (\text{S1})$$

### I. High-salt Dissolution Assay

High-salt dissolution was used to qualitatively assess condensate glassiness, complementing FRAP results. A 1 M KCl solution, at 10 times the sample volume, was added to the sample well containing FUS condensates aged for different time periods (1 hour, 24 hours, and 48 hours) at room temperature (23 °C). Confocal fluorescence images were acquired immediately after the addition of 1 M KCl to initiate condensate dissolution. To quantify condensate area over time, we used the maximum intensity projection of each confocal z-stack. Condensate regions were segmented using Otsu's thresholding method to generate binary masks for each frame. Since dissolving condensates gradually lose their circular morphology, we quantified area by summing the number of foreground (non-zero) pixels within the thresholded masks. Areas were normalized to the first-frame value for each time series.

### II. IMAGE ANALYSIS

#### A. Proteostat fluorescence intensity quantification

To determine proteostat intensity in a fiber-growing series, the summed projection of a confocal z-stack is used. For the graph in Fig. 1e, we integrate the total fluorescence intensity at each time point, $I(t)$. We subsequently normalize the data using the minimum and maximum intensity, $I_0$ and $I_{\max}$ respectively.

For the graphs in Fig. 4c, we were interested in quantifying the percentage increase of fibers from the initial value. We reasoned that the intensity of the fiber seed was a good measure of a typical fiber intensity while the initial dilute phase signal was a good measure of background. We use Otsu's method to threshold our summed image and identify high-intensity regions as seeds in the initial time stack. We thereafter define the value, $\bar{I}_{prot} = \Sigma I_{x,y}(t) - \langle I_{xy,bkgrnd} \rangle$. This is the pixel-by-pixel intensity, $I_{x,y}$ corrected for the average background intensity, $\langle I_{xy,bkgrnd} \rangle$. We determine $\langle I_{xy,bkgrnd} \rangle$ as the average intensity of pixels not colocalized with the high-intensity regions identified as seeds only in the initial time point. We normalize $\bar{I}_{prot}$ by its value in the first time point, $\bar{I}_{prot,0}$. To determine the plateau values at the end of the intensity curves, we took the mean value for the final 15 points along with the standard deviation. For the initial gradient of the curves we fit the first 5 points to a straight line using a least squares method, with the error quoted being the square root of the variance of the fit.

### B. Quantification of GFP intensity to obtain condensate area and radius over time

For the quantification of condensate area in Fig. 1e we take our imaging series of the GFP and proteostat signals of samples at early age (liquid-like condensates) imaged using confocal microscopy. At each time point we create a maximum projection of the confocal stack resulting in a single image using FIJI. Using custom scripts in MATLAB, we threshold the proteostat signal similar to the procedure described in section II A. We also threshold the GFP signal at each time point using Otsu’s method to identify high intensity regions corresponding to condensates and fibers. To measure the area of the condensates specifically, we remove any regions which overlap with a high signal in the proteost image using the thresholded proteostat data. To each of the condensate regions, the area of the condensate was estimated. The value $A(t)$ is defined as the sum of all condensate regions at each time point, $t$. In the graph shown in Fig. 1e, the value is normalized to the initial area, $A_0 = A(t = 0)$. For the condensate shown in Fig. 1f, blue dotted line, the data are treated similarly with custom Python scripts, the radius is determined from the condensate area of the outlined condensate and plotted in Fig. 1g.

To analyze the radius of condensates over time, we take our imaging series of from the GFP signal of samples with incubation times of $t_w = 1$ h, $t_w = 24$ h, and $t_w = 48$ h imaged using confocal microscopy. At each time point we create a maximum projection of the confocal stack resulting in a single image using FIJI. Using custom scripts in MATLAB we measure the radius of representative condensates at each time point. To ensure that our measurement does not suffer from apparent changes in radius owing to coalescence events or deformation from fiber growth, we identify and crop out condensates that do not move, coalesce or merge during the entire time course of observation. Briefly, the radius is obtained using the Otsu method to threshold the image. The major and minor axis of each thresholded condensate using the MATLAB function `regionprops`. We confirm the major and minor axis do not differ more than 10% and subsequently use the major axis as a measure of condensate radius. We average all radii at each time point for each sample. This average defines $R(t)$. For the graph in Fig. 4, the average radius, $R(t)$ is also scaled by its initial value, $R_0 = R(t = 0)$.

We use our theoretical prediction of the scaling of $R(t)$ in either the diffusion-limited regime or the interface limited region to fit the curves using the Python function 'curve fit' which utilizes a least squares algorithm. Fitting parameters are reported in the main manuscript and caption.

#### C. Estimation of fraction of fiber growth arising from conversion of condensed phase protein directly compared to the conversion from dilute phase protein

The volume of condensates was measured by first thresholding the GFP channel in ImageJ, with watershedding applied if necessary to distinguish individual condensate. A Python script then fit a bounding rectangle to each condensate using OpenCV [39], providing two radii for every condensate at each time point, one for the major axis, $r_1$ and one for the minor axis, $r_2$, of an ellipse. Assuming ellipsoidal geometry, condensate volumes were calculated from these radii, using the minor axis radius for the height of the condensate: $V = \frac{4}{3}\pi r(t)_1 r(t)_2^2$.

To classify condensates, the proteostat channel was thresholded in ImageJ. Condensates overlapping with the thresholded proteostat signal were labeled as colocalized condensates (labelled as red in Extended Data Fig. 3b), while non-overlapping condensates were classified as non-colocalized condensates (labelled as blue in Extended Data Fig. 3b). The total volume of colocalized and non-colocalized condensates was obtained by summing their respective volumes.

### III. THEORETICAL FRAMEWORK

This section details the theoretical framework used to describe the shrinkage of a phase-separated condensate and the associated formation of fibers. In our model, a spherical FUS condensate of radius $R(t)$ is centered within a system whose outer boundary, representing the fiber's position, is at a distance $R_F$. We denote the FUS monomer concentration profile in the external phase as $\phi(\underline{r}, t)$. The concentrations immediately inside and outside the droplet's phase boundary (at $|\underline{r}| = R$) are defined as $\phi_{\text{in}}$ and $\phi_{\text{out}}$, respectively. We assume that the interior of the droplet is well mixed so that the concentration inside the droplet is a constant ($\phi_{\text{in}}$), and thus there is no diffusional flux. At the outer boundary ($|\underline{r}| = R_F$), the concentration is $\phi_F$, which gives rise to a flux density $\underline{j}_F$. While the theory is developed for a general $\phi_F$, all results presented in the main text assume a perfect sink where $\phi_F = 0$.

The fundamental principle of our model is that the time evolution of the FUS monomer concentration in the dilute phase ($R(t) \leq |\underline{r}| \leq R_F$) is governed by the diffusion equation:

$$\partial_t \phi(\underline{r}, t) = D_{\text{out}} \nabla^2 \phi(\underline{r}, t), \quad (\text{S2})$$

where $D_{\text{out}}$ is the diffusion coefficient outside droplet. This equation must be solved subject to boundary conditions at the system's interfaces. At the outer boundary and the moving droplet surface, we can define the boundary condition as:

$$\phi(R_F) = \phi_F, \quad \phi(R) = \phi_{\text{out}}. \quad (\text{S3})$$

The system is closed by considering the dynamics of the droplet boundary itself. For a non-equilibrium shrinking droplet, the flux of FUS monomer across the phase boundary, $J$, can be expressed as a linear function of the thermodynamic driving force [35]:

$$J = \Gamma (\phi_{\text{in}} - P \phi_{\text{out}}). \quad (\text{S4})$$

Here, $P$ is the equilibrium partition coefficient ($P = \phi_{\text{in,eq}}/\phi_{\text{out,eq}}$) and $\Gamma$ is the interfacial conductance. Further, the flux balance condition in the co-moving frame of droplet boundary leads to

$$J = -\phi_{\text{in}} \dot{R} = j_{\text{out}} - \phi_{\text{out}} \dot{R}. \quad (\text{S5})$$

There is no flux inside the droplet ($j_{\text{in}} = 0$) in Eq. (S5) due to the well-mixed concentration inside the droplet.

Solving the full moving-boundary problem described above is complex. For this work, we make two key simplifying assumptions. First, we assume the system is spherically symmetric, reducing the problem to a single spatial dimension, $r$. Second, we assume that the diffusion of solute in the external phase is much faster than the rate at which the droplet radius changes. This allows us to make the quasi-static approximation, where the concentration profile $\phi(r)$ is assumed to instantaneously adapt to the current radius $R(t)$. In this limit, the time-derivative term in the diffusion equation vanishes, $\partial_t \phi = 0$, and the governing equation simplifies to the Laplace equation:

$$\nabla^2 \phi(r) = 0 \quad (R \leq r \leq R_F). \quad (\text{S6})$$

In the following sections, we derive the solution for this simplified, quasi-static problem. First, in section III A, we solve the Laplace equation to determine the external concentration profile and the droplet shrinkage rate, $\dot{R}$. In the successive section III B, we use this solution to calculate the rate of accumulated fibers at the sink. Finally, in section III C, we derive the limiting expressions for the condensate radius as a function of time for the diffusion-limited and interfacial-limited regimes.

#### A. The model of condensate's shrinkage with fiber formation

We describe the concentration profile outside of the droplet using Laplace's equation. We assume that the outside the droplet ($R \leq r \leq R_F$) the concentration is quasi-steady and spherically symmetric in three dimension, thus

$$\nabla^2 \phi = \frac{1}{r^2} \frac{\partial}{\partial r} \left( r^2 \frac{\partial \phi}{\partial r} \right) = 0 \quad (\text{S7})$$

leading to

$$\phi(r) = C_1 + \frac{C_2}{r}, \quad (\text{S8})$$

where $C_1$ and $C_2$ are the constants determined by the boundary conditions. Using the boundary condition, Eq. (S3), the solution of the Laplace equation is given as

$$\phi(r) = \phi_F + (\phi_{\text{out}} - \phi_F) \frac{R}{r} \frac{R_F - r}{R_F - R}, \quad (R \leq r \leq R_F). \quad (\text{S9})$$

Differentiate Eq. (S9) and evaluate at $r = R$ leads to

$$\left. \frac{d\phi}{dr} \right|_{r=R} = -\frac{\phi_{\text{out}} - \phi_F}{R(1 - R/R_F)}. \quad (\text{S10})$$

With Fick's law, the material flux outside the droplet is (outward positive),

$$j_{\text{out}} = \frac{D_{\text{out}} (\phi_{\text{out}} - \phi_F)}{R(1 - R/R_F)}. \quad (\text{S11})$$

Combining this with $\dot{R} = -j_{\text{out}}/(\phi_{\text{in}} - \phi_{\text{out}})$ from (S5), we obtain

$$\dot{R} = -\frac{D_{\text{out}} (\phi_{\text{out}} - \phi_F)}{R(1 - R/R_F) (\phi_{\text{in}} - \phi_{\text{out}})}. \quad (\text{S12})$$

Equating Eq (S12) with $\dot{R} = -\Gamma(\phi_{\text{in}} - P\phi_{\text{out}})/\phi_{\text{in}}$ from Eq. (S4)-(S5), we obtain quadratic in $\phi_{\text{out}}$:

$$\Gamma R (\phi_{\text{in}} - \phi_{\text{out}}) (\phi_{\text{in}} - P\phi_{\text{out}}) = D_{\text{out}} \phi_{\text{in}} \frac{\phi_{\text{out}} - \phi_F}{1 - R/R_F}. \quad (\text{S13})$$

Introducing the dimension-less quantity

$$x(R) = \frac{\Gamma R(1 - R/R_F)}{D_{\text{out}}}, \quad (\text{S14})$$

Eq. (S13) leads to

$$x(\phi_{\text{in}} - \phi_{\text{out}})(\phi_{\text{in}} - P\phi_{\text{out}}) = \phi_{\text{in}}(\phi_{\text{out}} - \phi_F). \quad (\text{S15})$$

The solution of the quadratic equations are

$$\phi_{\text{out}}^{\pm}(R) = \frac{-B \pm \sqrt{B^2 - 4AC}}{2A}, \quad (\text{S16})$$

with

$$A = xP; \quad B = -\phi_{\text{in}}(x(1 + P) + 1); \quad C = x\phi_{\text{in}}^2 + \phi_F\phi_{\text{in}}. \quad (\text{S17})$$

The $\phi_{\text{out}}^-(R)$ is the physical solution for shrinking droplet. We obtain ordinary differential equation for $R(t)$ by substitute the solution into Eq. (S12):

$$\dot{R} = -\frac{D_{\text{out}}(\phi_{\text{out}}(R) - \phi_F)}{R(1 - R/R_F)(\phi_{\text{in}} - \phi_{\text{out}}(R))}. \quad (\text{S18})$$

### B. Fiber formation

The point $r = R_F$ is a sink which represents fiber formation. By considering the flux at this point, we can estimate the total protein concentration lost to fiber formation. We differentiate Eq. (S9) with respect to $r$ and evaluate at $r = R_F$ to obtain,

$$\left. \frac{d\phi}{dr} \right|_{r=R_F} = -\frac{\phi_{\text{out}} - \phi_F}{R_F(R_F/R - 1)}. \quad (\text{S19})$$

With Fick's law (outward positive),

$$j_F = \frac{D_{\text{out}}(\phi_{\text{out}} - \phi_F)}{R_F(R_F/R - 1)}. \quad (\text{S20})$$

Therefore the total material volume per unit time that goes to fiber is

$$J_F = 4\pi R_F^2 j_F = 4\pi \frac{R_F D_{\text{out}}(\phi_{\text{out}} - \phi_F)}{(R_F/R - 1)}. \quad (\text{S21})$$

In the main text, Fig. 3, we report the total concentration of fiber which is obtained by integrating Eq. (S21) in time.

#### C. Limiting regimes at large and small interfacial conductivity

In this section, we obtain the expressions for the condensate radius in the interface-limited and diffusion-limited regime, $R(t) = -\Gamma t + R_0$ and $R(t) = \sqrt{-at + R_0^2}$, respectively.

##### 1. Interface-Limited Regime (Small $\Gamma$)

This regime occurs when the interfacial transfer rate is much slower than the characteristic rate of diffusion: $\Gamma/(D_{\text{out}}/R) \rightarrow 0$. Consequently, the dimensionless number $x \rightarrow 0$. As $x \rightarrow 0$, the left-hand side of Eq. (S15) approaches zero. Assuming $\phi_{\text{in}} \neq 0$, this implies:

$$\phi_{\text{out}} = \phi_F. \quad (\text{S22})$$

In this limit, diffusion is fast enough to maintain the far-field concentration $\phi_F$ at the droplet interface, and the process is limited by the slow kinetics of solute transfer across the interface.

The shrinkage rate is governed by the interfacial kinetic law, $J = \Gamma(\phi_{\text{in}} - P\phi_{\text{out}})$. The co-moving flux balance also gives $J = -\phi_{\text{in}}\dot{R}$. Equating these and using $\phi_{\text{out}} = \phi_F$:

$$\dot{R} = -\frac{\Gamma}{\phi_{\text{in}}}(\phi_{\text{in}} - P\phi_F). \quad (\text{S23})$$

Let $k = \frac{\Gamma}{\phi_{\text{in}}}(\phi_{\text{in}} - P\phi_F)$. For the droplet to shrink, $k > 0$, requiring $\phi_{\text{in}} > P\phi_F$. The rate of shrinkage is:

$$\dot{R} = -k. \quad (\text{S24})$$

If $\Gamma$ (and other parameters) are constant, $\dot{R}$ is constant. Therefore we obtain

$$R = -kt + R_0. \quad (\text{S25})$$

If we set $\phi_F = 0$, which is reasonable at fiber position, then we simply obtain

$$R = -\Gamma t + R_0. \quad (\text{S26})$$

Eq. (S29) is the expression for the interface-limited regime used in the main text.

##### 2. Diffusion-Limited Regime (Large $\Gamma$)

This regime occurs when the interfacial transfer rate of the proteins is much faster than the characteristic rate of diffusion: $\Gamma/(D_{\text{out}}/R) \rightarrow \infty$. Consequently, the dimensionless number $x \rightarrow \infty$.

As $x \rightarrow \infty$ in Eq. (S15), for the right-hand side to remain finite, we must have:

$$(\phi_{\text{in}} - \phi_{\text{out}})(\phi_{\text{in}} - P\phi_{\text{out}}) = 0. \quad (\text{S27})$$

This implies either $\phi_{\text{out}} = \phi_{\text{in}}$ or $\phi_{\text{out}} = \phi_{\text{in}}/P$. If $\phi_{\text{out}} = \phi_{\text{in}}$, the denominator $(\phi_{\text{in}} - \phi_{\text{out}})$ in Eq. (S18) would approach zero, leading to an unphysically infinite $\dot{R}$. The physically relevant solution for a diffusion-controlled process with fast interfacial kinetics is that local thermodynamic equilibrium is established at the interface, meaning the kinetic driving force $\phi_{\text{in}} - P\phi_{\text{out}}$ vanishes. Thus:

$$\phi_{\text{out}} = \frac{\phi_{\text{in}}}{P}. \quad (\text{S28})$$

Substituting Eq. (S28) into the expression for $\dot{R}$ [Eq. (S18)]:

$$\dot{R} = -\frac{D_{\text{out}}(\phi_{\text{in}}/P - \phi_F)}{R(1 - R/R_F)(\phi_{\text{in}} - \phi_{\text{in}}/P)} = -\frac{D_{\text{out}}}{R(1 - R/R_F)} \left( \frac{\phi_{\text{in}}/P - \phi_F}{\phi_{\text{in}}(1 - 1/P)} \right). \quad (\text{S29})$$

Let $S = \frac{\phi_{\text{in}}/P - \phi_F}{\phi_{\text{in}}(1 - 1/P)} = \frac{\phi_{\text{in}} - P\phi_F}{\phi_{\text{in}}(P - 1)}$. For shrinking droplets, $S > 0$. Given $P > 1$, this requires $\phi_{\text{in}} > P\phi_F$. The rate of shrinkage is

$$\dot{R} = -\frac{D_{\text{out}}S}{R(1 - R/R_F)}. \quad (\text{S30})$$

Rearranging Eq. (S30):

$$\left( R - \frac{R^2}{R_F} \right) dR = -D_{\text{out}}S dt. \quad (\text{S31})$$

Integrating from $R_0$ at $t = 0$ to $R(t)$ at time $t$:

$$\int_{R_0}^{R(t)} \left( R' - \frac{R'^2}{R_F} \right) dR' = \int_0^t -D_{\text{out}}S dt'. \quad (\text{S32})$$

This gives the implicit relationship for $R(t)$:

$$\left( \frac{R_0^2}{2} - \frac{R(t)^2}{2} \right) - \left( \frac{R_0^3}{3R_F} - \frac{R(t)^3}{3R_F} \right) = D_{\text{out}}St. \quad (\text{S33})$$

If the outer boundary is very far away, or the droplet is much smaller than $R_F$, Eq. (S33) simplifies to

$$R(t)^2 = -(2D_{\text{out}}S)t + R_0^2. \quad (\text{S34})$$

If we set the $\phi_F = 0$, which is reasonable at the fiber position, then we obtain

$$R(t)^2 = -at + R_0^2; \quad a = 2D_{\text{out}}/(P - 1). \quad (\text{S35})$$

Eq. (S35) is the expression for the diffusion-limited regime used in the main text.

| Parameter | Meaning | Value | Unit |
| --- | --- | --- | --- |
| $\phi_{\text{in}}$ | Inside concentration | 1 | *none* |
| $\phi_F$ | concentration at fiber position | 0 | *none* |
| $P$ | Partition coefficient | $3.0 \times 10^5$ | *none* |
| $D_{\text{out}}$ | Diffusion coefficient outside | 30 | $\mu\text{m}^2/\text{s}$ |
| $R_0$ | Initial radius of droplet | 3 | $\mu\text{m}$ |
| $R_F$ | Fiber position | 100 | $\mu\text{m}$ |

TABLE I: Parameters used in Fig. 3 in the main text.
